# A zebrafish seizure model of *cblX* syndrome reveals a dose-dependent refractory response to mTor inhibition

**DOI:** 10.1101/2025.11.14.688058

**Authors:** Claudia B. Gil, David Paz, Briana E. Pinales, Victoria L. Castro, Claire E. Perucho, Annalise Gonzales, Giulio Francia, Sepiso K. Masenga, Antentor Hinton, Anita M. Quintana

## Abstract

Mutations in the transcriptional cofactor HCFC1 cause methylmalonic aciduria and homocystinemia, cblX type (*cblX*) (MIM#309541), non-syndromic X-linked intellectual disability (XLID), and focal epilepsy. Zebrafish studies have revealed increased activation of the Akt/mTor signaling pathway after mutation of *hcfc1a*, one ortholog of *HCFC1*. mTOR hyperactivation is linked to seizures and its inhibition alleviates epilepsy in other preclinical models. We hypothesized that mTor overactivity in *hcfc1a* mutant zebrafish increases seizure susceptibility and/or severity. We employed a two-concentration model of the seizure inducing agent, pentylenetetrazol (PTZ), with or without pretreatment of the mTor inhibitor, torin1. Mutation of *hcfc1a* did not increase seizure susceptibility at sub-optimal concentrations of PTZ and the pharmaceutical inhibition of mTor reduced seizure severity when utilized at a dose of 250nM. Higher doses of torin1 treatment exacerbated seizure response in mutant larvae but not in wildtype siblings. These data suggest that in an *hcfc1a* deficient background, moderate inhibition of mTor signaling may partially alleviate seizure phenotypes, however, over-inhibition of the pathway causes a refractory response to PTZ. Collectively, we present a model that can be used to test dose response and for the development of combinatorial treatment approaches in a high throughput manner.

## Introduction

*HCFC1* encodes a multi-domain transcriptional co-factor that regulates the expression of more than 5000 genes [1]. Mutations in the kelch protein interaction domain of HCFC1 cause methylmalonic aciduria with homocystinemia, type cblX (*cblX*), a multiple congenital anomaly syndrome characterized by abnormal vitamin B12 metabolism, craniofacial dysmorphia, neurodevelopmental defects, failure to thrive, and intractable epilepsy. Mutations in other domains of HCFC1 cause non-syndromic X-linked intellectual disability (XLID) or focal epilepsy [2–6]. The patient phenotypes associated with pathogenic variants of HCFC1 suggest that the protein is essential for neural development and its dysfunction may increase susceptibility and the severity of seizure phenotypes.

Functional analysis has demonstrated that HCFC1 is required to maintain the number and proliferation of neural precursor cells (NPCs) and neurite outgrowth [2,3]. Subsequent studies in zebrafish validated the increased number and proliferation of NPCs through morpholino mediated knockdown and germline mutation of the *hcfc1a* and *hcfc1b* paralogs, respectively [7,8]. Nonsense mutation of murine *Hcfc1* in a sub-population of NPCs resulted in increased cell death and reduced GABAergic cells [9], the primary inhibitory cells of the central nervous system, whose dysfunction has been implicated in epilepsy and seizure. A genetic knock-in allele of *cblX* syndrome was created in mice but brain phenotypes were not comprehensively studied [10]. Consequently, the cellular phenotypes and molecular mechanisms regulating brain, seizure, and behavioral phenotypes in *cblX* syndrome are not completely understood. Moreover, there is currently no validated model or paradigm optimized to study seizure phenotypes in *cblX*.

Nonsense mutation of the zebrafish *hcfc1a* gene (*hcfc1a^co60/+^*) resulted in increased expression of the *asxl1* gene [8]. *Asxl1* encodes a polycomb repressor protein with functions in the nucleus and cytoplasm. ASXL1 in the cytoplasm leads to hyperphosphorylation of AKT kinase in mouse embryonic fibroblasts [11]. AKT kinase regulates cell growth, proliferation, and survival. Mutations in the AKT pathway are associated with epileptic phenotypes [12]. Interestingly, inhibition of Akt (zebrafish) phosphorylation and activation restores NPC overproduction in the *hcfc1a^co60/+^* haploinsufficient zebrafish, providing evidence that Akt and its downstream pathways underly some neurodevelopmental phenotypes observed in the mutant larvae. Subsequent experiments demonstrated hyper-activation of Akt and mTor (mammalian target of rapamycin) in the *hcfc1a^co60/+^* zebrafish [13]. Interestingly, hyperactivation of AKT/mTOR signaling has been extensively shown to underly major seizure disorders, such as Tuberous Sclerosis Complex [14–17]. Based on these data, we hypothesized that mutation of *hcfc1a* leads to increased seizure susceptibility and severity due to hyperactivation of mTor.

In utero seizures have been detected in humans with mutations in *HCFC1* [18]. Thus, we opted to monitor seizure phenotypes in the larval zebrafish after exposure to pentylenetetrazol, a GABA receptor antagonist, established to cause seizures in multiple models systems [19–22]. We selected the *hcfc1a^co60/+^* genotype for our studies because these zebrafish carry a nonsense mutation in the kelch domain, are heterozygous viable, and have increased numbers of NPCs with hyperactive mTor signaling [8,13]. We exposed larval zebrafish (*hcfc1a^co60/+^* offspring) to established seizure inducing concentrations of PTZ and sub-optimal doses (a dose that has no behavioral effect on wildtype animals). The sub-optimal dose did not increase sensitivity to seizure induction in larvae with the *hcfc1a^co60/+^* allele. Wildtype siblings had a reduced response to PTZ when pretreated with mTor inhibitors and the efficacy of pharmaceutical inhibition improved with higher concentrations of mTor inhibition. In contrast, the efficacy of pretreatment in mutant larvae was dose dependent, with lower concentrations showing improved efficacy and higher concentrations exacerbating the seizure-like phenotypes according to the metrics evaluated. A substantial amount of mTor remained active in mutant larvae despite pretreatment with an mTor inhibitor, substantiating dysregulation of mTor in mutants. These data indicate that while mutation of *hcfc1a* and hyperactivation of mTor do not increase seizure susceptibility, the manipulation of mTor, in the context of *hcfc1a* mutation, can contribute to intervention efficacy and can promote a refractory and deleterious response at high doses, which show improved efficacy in wildtype animals. Follow up combinatorial therapies with a full panel of inhibitors, multiple treatment windows, and additional dose response curves are warranted. These follow up studies can be tested in a high throughput manner when used in combination with our current model system and seizure paradigm.

## Materials and Methods

### Zebrafish and genotyping

For the following experiments, embryos are produced from natural spawning of the following lines: *hcfc1a^co60/+^*, AB, Tupfel long fin, or AB crossed with Tupfel long fin. The *hcfc1a^co60/+^* is maintained in a background that carries AB and Tupfel long fin genetic inheritance. Embryos are maintained in E3 media (embryo medium) at 28^◦^ with a 14/10 light: dark cycle. All procedures were approved by the Institutional Animal Care and Use Committee at the University of Texas at El Paso, protocol number 811869-5. Methods for euthanasia and anesthesia were performed according to guidelines from the 2020 American Veterinary Medical Association guidebook.

All adults and larvae were genotyped using polymerase chain reaction (PCR). Tissue was lysed in 50 millimolar (mM) sodium hydroxide (Fisher Scientific) for 10 minutes at 95^◦^ Celsius and pH adjusted with 500 millimolar (mM) Tris-HCl. Allele-specific primer pairs were used for each allele. These primers bind to and amplify the mutated allele but do not amplify the wild-type allele (5). For the *hcfc1a*^co60/+^ allele, the primers specific to the mutated allele were FWD: CCAGTTCGCCTTTTTGTTGT and REV: ACGGGTGGTATGAACCACTGGC, each used at a final concentration of 0.5mM. PCR annealing was executed at 64° and 30-35 cycles were performed. GoTaq Green (1X) was used for all genotyping (Fisher Scientific).

### Zebrabox Behavioral Analysis

Behavioral analysis was performed using the Zebrabox technology (ViewPoint Behavioral Technology, Montreal Canada). Analysis was performed at 5 days post-fertilization (dpf). The presence of a swim bladder was confirmed prior to the analysis. Larvae were individually placed into a 48-well plate with E3 media. All larvae were acclimated in dark conditions for 1 hour before data collection. The behavioral assay lasted for a total duration of 10 minutes in the dark. After acclimation, behavior was recorded for 5 minutes in the dark to obtain a baseline measurement. Parameters were developed based on previous studies [23,24] except for black detection used instead of the transparent setting and the threshold was set to 25 to provide improved detection in dark conditions. The Zebrabox reports the duration (s) and distance of swimming (mm).

Swimming measurements are split into small and large. The following parameters: total distance, large distance, small distance, speed, large duration, small duration, large count, and small count, were analyzed to determine changes in locomotor activity and seizure-like behavior. The raw data was recorded on an excel sheet along with a video file. Speed and total distance were manually calculated as described [23]. Total distance is the sum of large and small distances and speed is calculated by dividing total distance by the sum of small and large duration parameters (mm/s). Trace patterns are color coded in Figure 1 to indicate speed.

**Figure 1:**
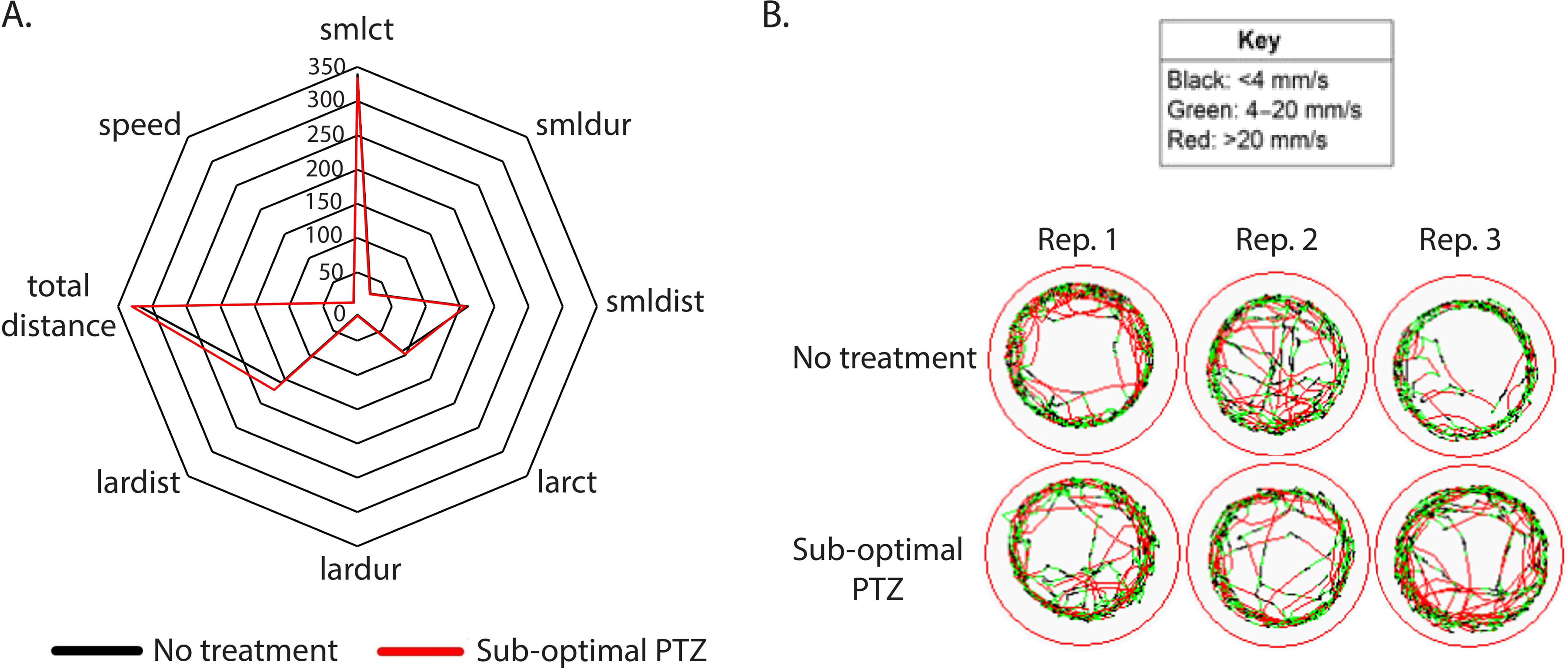
Empirical derivation of a sub-optimal concentration of PTZ. (A&B) Wildtype larvae (5 days post fertilization) were individually placed in a 48 well dish, acclimated for 1 hour to the ZebraBox environment, monitored 5 minutes for baseline behavior, and exposed to pentylenetetrazol (0.001pM). Radar plot of each parameter analyzed (A) and representative trace patterns are shown (B). N=48/group. Abbreviations: Rep: Replicate

### Torin1 and PTZ inhibition

A sub-optimal concentration of PTZ was derived empirically by performing a concentration gradient and monitoring seizure like behaviors (locomotion, distance, time, and speed). The gradient was performed using wildtype larvae at 5 dpf. Larvae were individually placed into a 48-well plate with 1,450µL of E3 media. After acclimation and a 5-minute baseline recording, PTZ was added to the following final concentrations 1µM, 10pM, 1pM, 0.1pM, 0.01pM and 0.001pM. Behavior was recorded for 5 minutes in the dark after exposure to PTZ. A sub-optimal dose of PTZ was determined and a total of 42 larvae were analyzed from multiple heterozygous outcrosses on different days, from different parents, raised to 5 dpf, analyzed for behavior, and subsequently genotyped. There were a total of 23 *hcfc1a*^co60/+^ larvae and 19 siblings. The following parameters: total distance, large distance, small distance, speed, large duration, small duration, large count, and small count, were analyzed to determine changes in locomotor activity and seizure-like behavior in response to the drug treatment. Seizure-like behavior was interpreted using the zebrafish behavioral catalog [25].

Torin1 is a selective inhibitor of mTOR (both complex 1 and 2) causing reduced cell growth and proliferation [26]. Western blot analysis was performed as previously described [13] to determine the appropriate concentration of torin1 required to reduce the level of phosphorylated S6 ribosomal protein in wildtype larvae. Based on these data, a concentration of 250nM of torin1 was deemed the minimal concentration to reduce phosphorylated S6 ribosomal protein, without resulting in abnormal development (gross morphological defects). Torin1 treatment was initiated at 24 hours post fertilization (hpf) to allow for early development. Torin1 was replaced daily for 4 days prior to behavioral analysis and treated larvae were challenged with PTZ at 5 dpf. To assess the effects of torin1 treatment on seizure severity, 1µM PTZ was used to induce seizure like behavior.

### Statistical Analysis

An ANOVA was used to determine the statistical differences between multiple groups followed by a post hoc T-test between individual groups. All data reported statistically significant has a p-value <0.05.

## Results

### Larval Response to Low Dose PTZ

We hypothesized that nonsense mutation of *hcfc1a* causes increased seizure susceptibility due to an underlying hyperactivation of mTor signaling [13,27]. To test this hypothesis, we first sought to identify a sub-optimal dose of the seizure inducing agent PTZ, by performing a concentration gradient on wildtype larvae at 5 dpf. Our goal was to find a concentration whereby wildtype larvae do not respond to PTZ, thus being sub-optimal in nature. We hypothesized that this sub-optimal concentration would elicit a heightened response in larvae with mutation in *hcfc1a* relative to their wildtype siblings, rendering the allele more susceptible to seizure-like behavior. We monitored seizure like behavior using ZebraBox technology after exposure to 1µM, 10pM, 1pM, 0.1pM, 0.01pM and 0.001pM PTZ. In Figure 1A, we generated a radar plot to create a visual summary of the multiple parameters that were tracked using Viewpoint automated technology. These parameters include total distance, large distance, small distance, speed, large duration, small duration, large count, and small count. Seizure-like behavior is described as whirlpool like behavior [28] with rapid movements that can be modeled by parameters such as increased small distance (bursts), increased time spent in rapid movements (small duration), and hyperlocomotion characterized by increased speed and total distance swam. As shown in Table 1; 1µM, 10pM, 1pM, and 0.1pM all induced a response after PTZ exposure, however to different degrees. For example, seven unique parameters were increased after exposure to 1µM PTZ. As we reduced the concentration of PTZ, we continued to observe considerable effects on the larvae, inducing behaviors indicative of seizure phenotypes such as hypermotility, increased movement, increased total distance, and increased time spent in small duration movements. The number of parameters significantly affected by exposure was reduced as the dose of PTZ was reduced, except for 10pM, which only led to the alteration of 1 parameter. In general, PTZ exposure increased movement and motility, however, at the 0.01pM concentration we observed reduced small count, small duration, small distance, large count, large duration, and total distance (Table 1). While it is counterintuitive that PTZ would elicit hypoactivity at any concentration, our observations at 0.01pM are consistent with previous studies that have shown that low concentrations of PTZ can have opposite effects relative to higher concentrations [29]. At the concentration of 0.001pM, we observed no response to PTZ (Figure 1A&B). Based on this data, we concluded that a concentration of 0.001pM of PTZ is a sub-optimal dose that has no seizure inducing activity in wildtype animals at 5 dpf.

**Table 1.** Empirical derivation of sub-optimal concentrations of PTZ and parameters monitored.

| PTZ | Small Count | Small Duration | Small Distance | Large Count | Large Duration | Large Distance | Total Distance (mm) | Speed | # of Parameters Affected |
| --- | --- | --- | --- | --- | --- | --- | --- | --- | --- |
| 1.0 μM | +59.17 | NC | +33.32 | +39.07 | +5.42 | +66.32 | +99.62 | +0.67 | 7 |
| 10 pM | NC | NC | NC | NC | NC | NC | NC | +0.83 | 1 |
| 1 pM | +194.93 | +18.73 | +116.85 | NC | NC | NC | +108.63 | -1.56 | 5 |
| 0.1 pM | +51.26 | NC | NC | +19.70 | +2.55 | NC | NC | NC | 3 |
| 0.01 pM | -49.65 | -4.92 | -32.97 | -22.48 | -2.62 | NC | -59.34 | NC | 6 |
| 0.001 pM | NC | NC | NC | NC | NC | NC | NC | NC | 0 |

NC: No significant change was identified. The values provided in the table indicate the increase (+) or decreased (-) response relative to untreated control larvae.

### Mutation of hcfc1a does not increase seizure susceptibility to low dose PTZ

The *hcfc1a^co60/+^* allele is a heterozygous viable, haploinsufficient nonsense mutation in the zebrafish *hcfc1a* gene, which has been characterized with abnormal neural development, behavioral deficits, and hyperactivated mTor [8,13]. *hcfc1a* is one conserved ortholog of *HCFC1* and therefore, represents a putative system to study the function of HCFC1 in brain disease. Given that mutation of *hcfc1a* results in hyperactive mTor, which has been associated with seizure, we hypothesized that treatment of the *hcfc1a^co60/+^* allele with a sub-optimal concentration of PTZ would elicit a seizure like response. We treated offspring of the *hcfc1a^co60/+^* allele with 0.001pM PTZ (sub-optimal) and developed a radar plot to analyze all parameters (total distance, large distance, small distance, speed, large duration, small duration, large count, and small count). As shown in Figure 2A&B, wildtype siblings and heterozygous carriers did not demonstrate significant differences in behavior for all parameters examined after treatment with PTZ (0.001pM). We analyzed heterozygous carriers because the allele is not homozygous viable. From the radar plot, we identified 4 parameters trending towards a significant response at sub-optimal doses of PTZ. For each of these parameters, we analyzed them independently using a post hoc T-test. These included small duration (smldur), small distance (smldis), large count (larct), and total distance. As shown in Figure 2C, smldur trended towards increasing but was not statistically significant. We did not observe any statistical change in smldist and larct (Figure 2D&E). Total distance was reduced (Figure 2F), but this change was not statistically significant supporting the conclusion that mutation of *hcfc1a* did not increase susceptibility to PTZ induced seizures.

**Figure 2.**
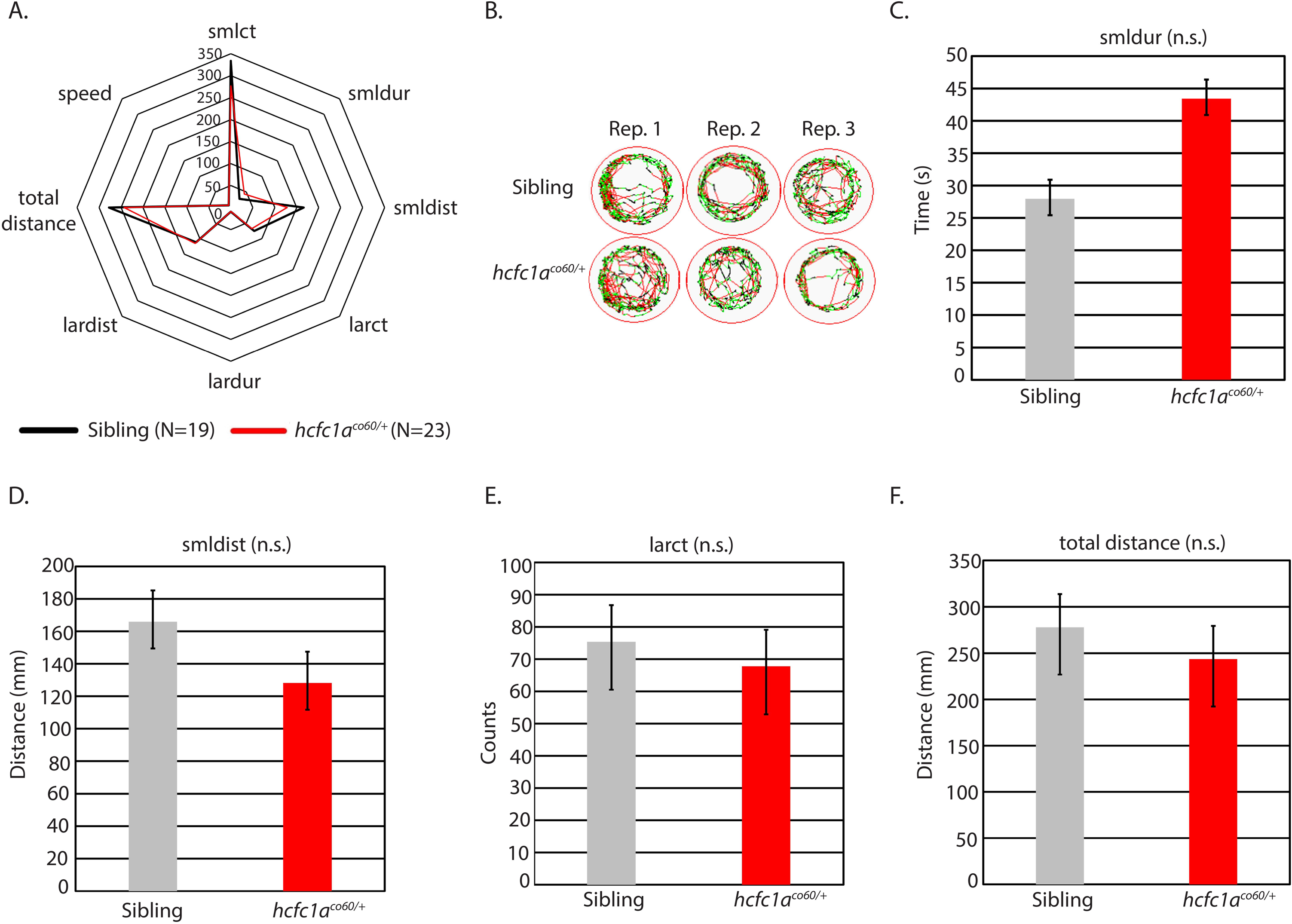
Mutation of *hcfc1a* does not increase seizure susceptibility to PTZ. (A) Offspring of the *hcfc1a^co60/+^* allele were treated with 0.001pM PTZ and assessed for behavioral patterns using the ZebraBox technology. Radar plot summarizes the behavior of sibling wildtype (black) and heterozygous mutants (red). (B) Representative trace patterns associated with behavior in (A). (C-F) Parameters from (A) for which there was a trending change in behavior were individually analyzed using a bar graph and T-test. Small duration (C), small distance (D), large count (E), and total distance (F) are shown. Error bars represent standard error of the mean. Sibling N=19; *hcfc1a^co60/+^* N=23

### Inhibition of mTor reduces seizure like behavior in wildtype animals

We sought to determine whether hyperactive mTor in the *hcfc1a^co60/+^* allele causes increased reactivity to seizure inducing concentrations of PTZ. We chose to treat larvae with 1µM PTZ because our dose gradient demonstrated 7/8 parameters increased after exposure (Table 1). Our first step was to inhibit mTor activity pharmacologically in wildtype larvae. We opted to inhibit mTor using torin1, a selective ATP competitive inhibitor of mTorc1 and mTorc2 complexes. We performed a concentration gradient (100-700nM) and monitored the level of phosphorylated S6 ribosomal protein (pS6) to determine efficacy of the treatment and validate inhibition of mTor (Figure 3A and Figure S1). We observed no decrease in pS6 at 100 and 200nM (Figure S1) but found decreased phosphorylation at concentrations equal to and higher than 250nM. Thus, we treated wildtype larvae with 250nM and 350nM concentrations of torin1 and then exposed pretreated larvae to seizure inducing concentrations of PTZ. As shown in Figure 3B and Table 2, treatment with 250nM torin1 reduced overall activity of wildtype larvae and caused a significant reduction in total distance swam and the number of times the larvae were detected in large movements (large count). Treatment with 350nM torin1 improved efficacy resulting in a statistically significant reduction in large count, large duration (time spent in large movements), and small count (number of times the larvae is detected in small movements). Total distance swam was also reduced but was not statistically significant (p=0.090). Thus, the increase in torin1 concentration was associated with a significant reduction in 3 parameters, whereas 250nM reduced only 2 parameters. These data demonstrate that pretreatment with an mTor inhibitor can effectively reduce seizure like behavior in zebrafish larvae.

**Figure 3.**
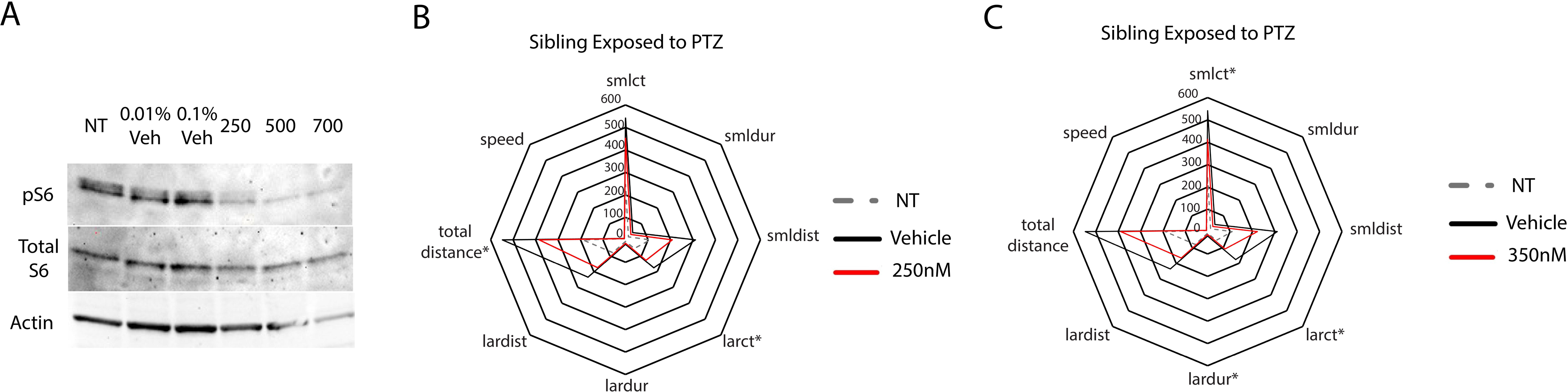
Optimization of torin1 treatment and PTZ response in wildtype animals. (A) Western blot with anti-phosphorylated ribosomal S6 protein, total S6 protein, and actin was performed on non-treated (NT), vehicle treated (different percentages), and the indicated concentrations of torin1 (250nM). For analysis in (A), 25 animals were harvested/group. Subsequently, animals were treated with 250nM (B) or 350nM (C) Torin 1 and then exposed to 1µM PTZ. For analysis shown in (B), animal numbers as follows: NT (N=22), Vehicle treated (N=20), and 250nM (N=19). Star plots were developed to indicate the effects of torin1 pretreatment. Asterisks indicate statistical significance (p<0.05). For analysis shown in (C) animal numbers are as follows: NT (N=22), vehicle treated (N=20), and 350 (N=18).

**Table 2.** Effectiveness of torin1 treatment on wildtype larvae.

| Torin1 | Small Count | Small Duration | Small Distance | Large Count | Large Duration | Large Distance | Total Distance (mm) | Speed | # of Parameters Affected |
| --- | --- | --- | --- | --- | --- | --- | --- | --- | --- |
| 250 nM | NC | NC | NC | -48.41 | NC | NC | -163.30 | NC | 2 |
| 350 nM | -127.81 | NC | NC | -67.52 | -7.35 | NC | NC | NC | 3 |

NC: No significant change was identified. The values provided in the table indicate the increase (+) or decreased (-) response relative to larvae treated with vehicle control and pentylenetetrazol.

### The effects of mTor inhibition on S6 phosphorylation

Next, we tested the effects of torin1 treatment on the *hcfc1a^co60/+^* allele. We hypothesized that a reduction in mTor signaling in mutant larvae would reduce the responsiveness to PTZ, given that mutation of *hcfc1a* results in hyperactive mTor signaling. To validate the effects of Torin1 in mutant animals, we performed western blot to detect pS6. Interestingly, treatment with vehicle control led to reduced levels of pS6 phosphorylation (Figure 4A). Consequently, we only compared behavior between the vehicle control groups and the torin1 treated groups. Torin1 completely abrogated pS6 in sibling wildtype, but torin1 treated mutants maintained a steady state level of pS6 relative to vehicle control treated mutants (Figure 4A). These data substantiate dysregulation of mTor signaling in mutant animals.

**Figure 4.**
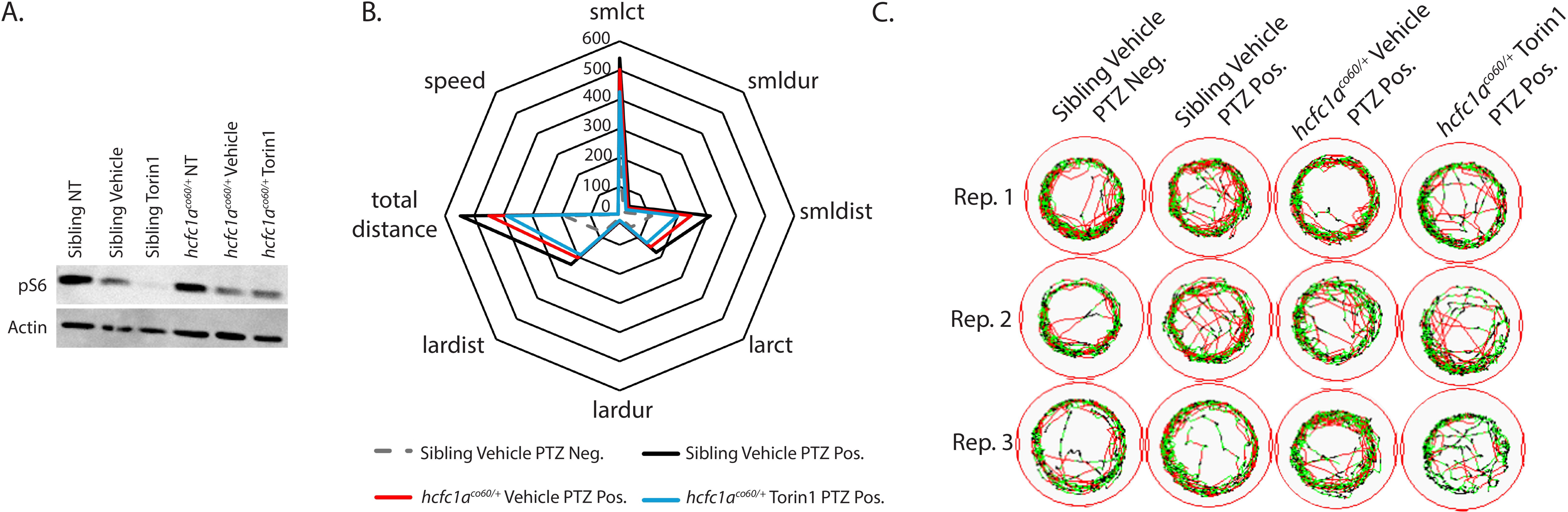
Effects of low dose torin1 treatment on pS6 and behavior. (A) Western blot was used to detect the phosphorylation of S6 ribosomal protein (pS6) in non-treated (NT), vehicle treated, or torin1 treated (250nM) wildtype siblings or *hcfc1a^co/60/+^* larvae. Actin is used as a loading control and additional ponceau stain was performed. N=14 larvae/group. (B) Radar plot analyzing 8 parameters collected using ZebraBox technology in larvae treated with vehicle or torin1 and exposed to 1µM PTZ (seizure inducing). Comparison is provided for animals with no PTZ exposure (neg) and those with PTZ exposure (pos). (C) Sample trace patterns from the animals in (B).

### Inhibition of mTor at 250nM reduces small/short behavioral parameters in mutant animals exposed to seizure inducing doses of PTZ

We hypothesized that inhibition of mTor in mutant larvae would reduce responsiveness to PTZ. We began our analysis by confirming the effects of 1µM PTZ on wildtype and mutant larvae at 5 dpf. Treatment with 1µM PTZ induced substantial seizure like phenotypes in sibling wildtype larvae treated with vehicle control (DMSO) (Figure 4B; black line relative to dashed gray line) as indicated by the fact that all parameters were significantly increased in wildtype siblings treated with vehicle control and 1µM PTZ. A similar response was observed in *hcfc1a^co60/+^* larvae treated with PTZ (Figure 4B and 5A-H). The mutant response to PTZ was reduced relative to wildtype but there was not a statistically significant change in the response between mutants and wildtypes in any parameter tested (Figure 4B; black vs red lines and Figure 5A-H). Representative trace patterns are shown in Figure 4C. Colored detection in Figure 4C monitors speed according to the key in Figure 1C.

**Figure 5.**
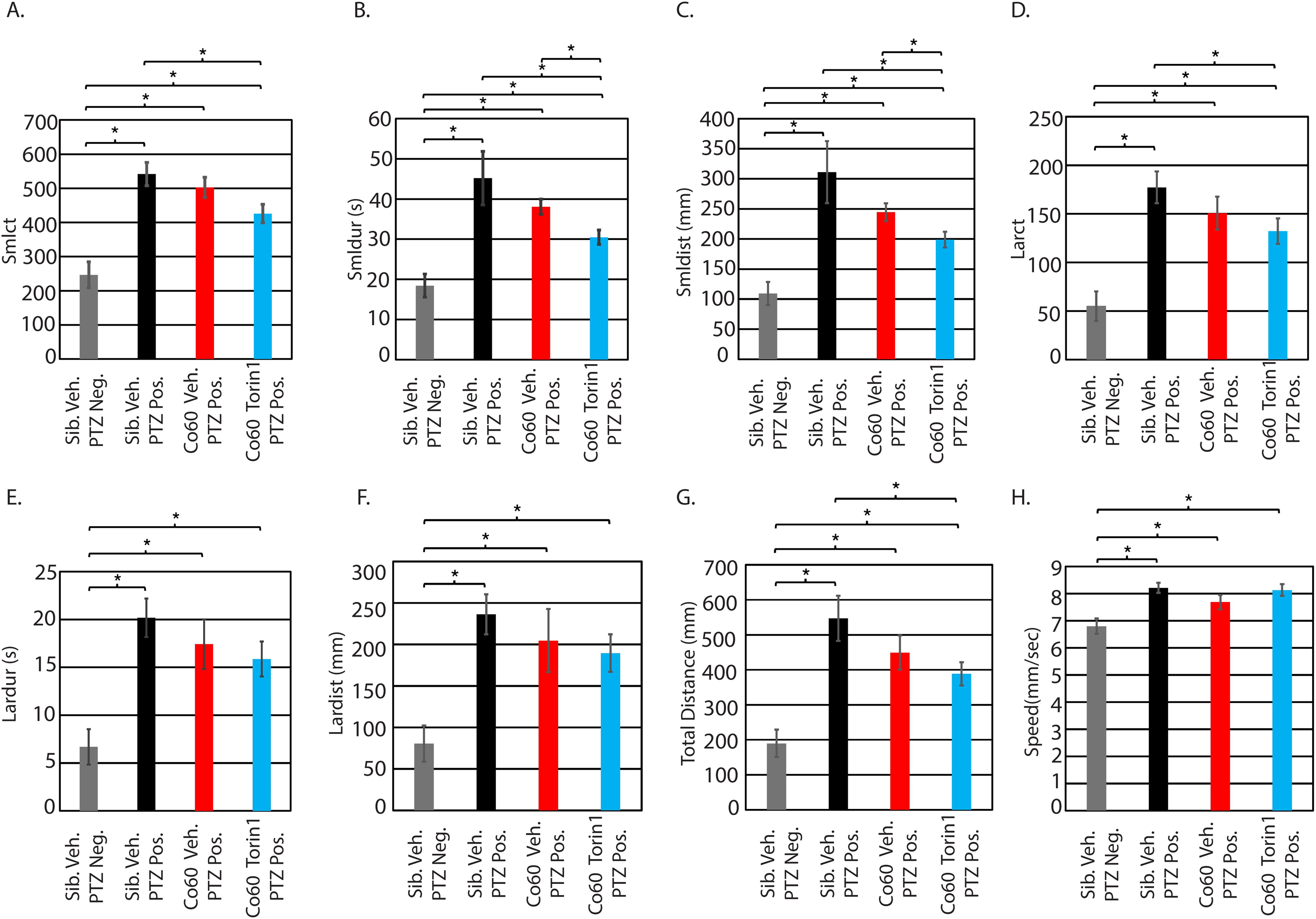
Inhibition of mTor reduces time spent in short distance and small duration movements. Parameters from Figure 4B were independently analyzed using ANOVA and post hoc T-test. Parameters include small count (A), small duration (B), small distance (C), large count (D), large duration (E), large distance (F), total distance (G), and speed (H). * p<0.05. Comparison is provided for animals with no PTZ exposure (neg) and those with PTZ exposure (pos). All error bars represent standard error of the mean. Sibling N=22; Vehicle *hcfc1a^co60/+^* N=28; torin1 *hcfc1a^co60/+^* N=28

We next analyzed the response of torin1 (250nM) treated mutant larvae relative to vehicle control mutant larvae, both exposed to 1µM PTZ. Initially, we detected a significant decrease in the response to PTZ in mutant larvae that were pretreated with torin1 (250nM) relative to vehicle treated sibling wildtype animals exposed to PTZ in the following parameters: smlct (Figure 5A), smldur (Figure 5B), smldist (Figure 5C), larct (Figure 5D), and total distance (Figure 5G). However, since vehicle treated mutants show a slightly reduced response to 1µM PTZ for all parameters relative to sibling vehicle exposed to PTZ (black bars vs red bars), we decided the best comparison to determine the positive effects of torin1 treatment (250nM) would be between vehicle control and torin1 treated (250nM) mutants, both exposed to 1µM PTZ. This approach allowed us to filter only the positive effects in mutant larvae. We detected a significant reduction in smldur and smldist between these groups (Figure 5B&C). These parameters are commonly associated with seizure-like behavior in zebrafish, as an increase in their values typically indicates heightened motility [25]. The observed decrease suggests that torin1 (250nM) partially mitigated seizure-like behavior (via reduction of smldur and smldist). However, these effects mirrored the efficacy of torin1 in wildtype larvae.

### Excessive inhibition of mTor in the context of hcfc1a mutation results in refractory epilepsy

As shown in Figure 3C, pretreatment of wildtype larvae with 350nM torin1 was more effective at reducing response to PTZ compared to 250nM. Treatment with 350nM reduced 3 parameters (lardur, larct, and smlct) whereas treatment with 250nM only reduced 2 parameters (larct and total distance). Based on these data, we hypothesized that pretreatment of mutant larvae with 350nM torin1 would be more efficacious than 250nM torin1. We predicted that higher levels of torin1 would consequently reduce more parameters. Consistent with our previous results (Figure 5A-H), wildtype and mutant larvae responded to PTZ, with mutant larvae showing a mild reduction in response relative to wildtype siblings. The reduction was not statistically significant in any parameter. In contrast to pretreatment with 250nM, mutant larvae treated with 350nM torin1 had an exacerbated response to PTZ characterized by increased smldur (Figure 6B) and smldist (Figure 6C) when compared to vehicle treated mutant larvae also stimulated with PTZ. In each parameter, mutants treated with PTZ and 350nM torin1 had a significant increase relative to sibling wildtype treated with vehicle control indicating that the pretreatment of 350nM torin1 was not effective at eliminating or alleviating the response to PTZ. Consequently, we conclude that pretreatment with 350nM torin1 does not improve the response of mutant larvae in any parameter (Figure 6A&D-H) and increases the response to PTZ for the smldur and smldist parameters (Figure 6B&C). These data contrast with the observed effects in wildtype larvae (Figure 3C) in which treatment with 350nM torin1 reduced activity in all parameters with small count, large duration, and large count demonstrating statistical significance.

**Figure 6.**
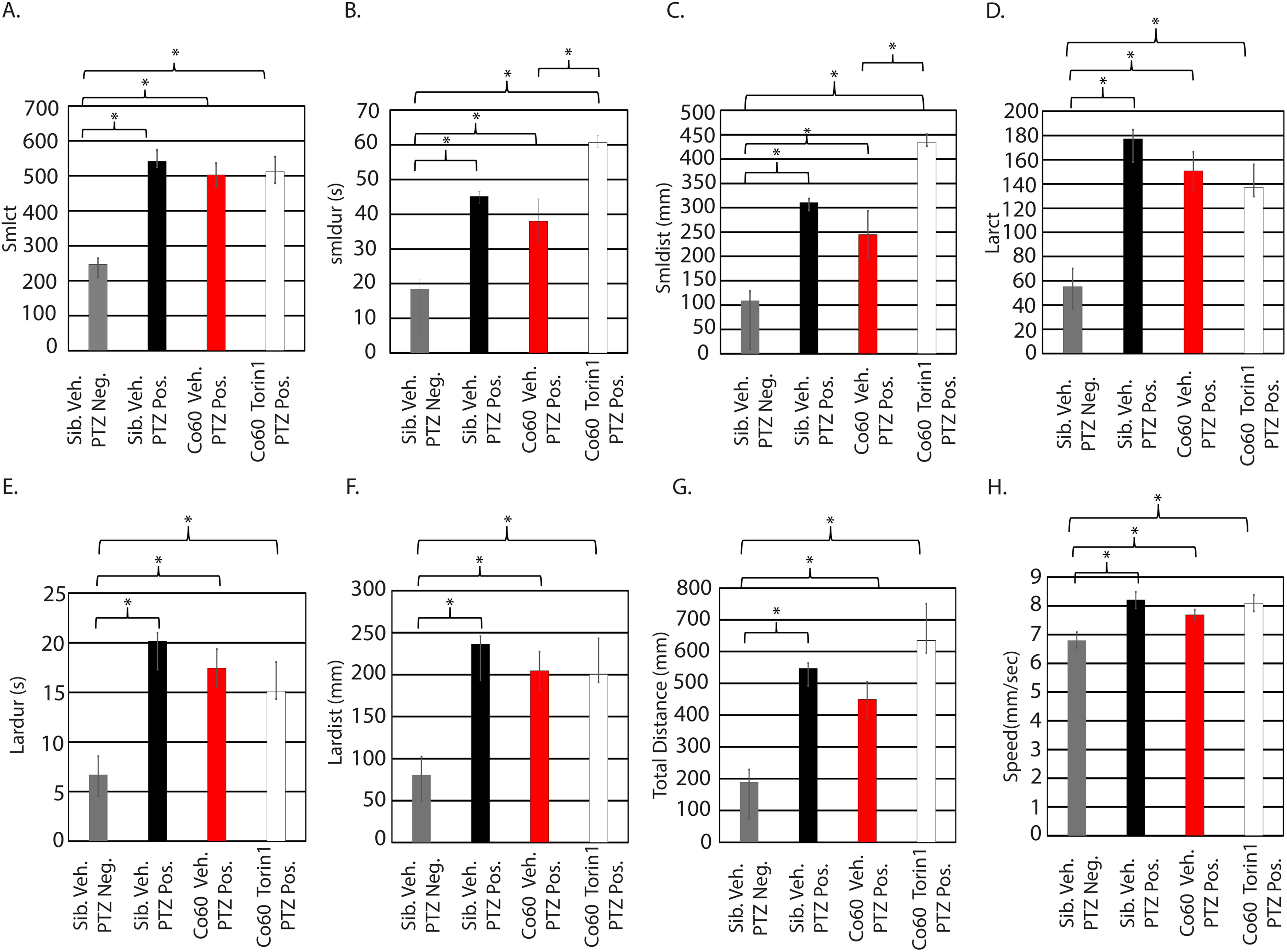
High dose inhibition of mTor exacerbates seizure like behavior in the *hcfc1a^co60/+^* allele. Zebrafish larvae were treated with 350nM torin1 and exposed to 1µM PTZ. Eight parameters were analyzed for each group: small count (A), small duration (B), small distance (C), large count (D), large duration (E), large distance (F), total distance (G), and speed (H). Numbers of animals per group are as follows: sib veh PTZ neg N=22, sib Veh PTZ Pos N=20, *hcfc1a^co60/+^* PTZ Pos N=28, *hcfc1a^co60/+^* torin1 PTZ Pos N=30

## Discussion

Our study demonstrates that mutation of *hcfc1a* does not increase seizure susceptibility. The pharmacological inhibition of mTor signaling can reduce the severity of PTZ-induced seizures in zebrafish harboring a nonsense mutation in *hcfc1a* but only in a dose dependent manner. While *hcfc1a* mutants did not exhibit increased sensitivity to sub-optimal PTZ doses, optimized mTor inhibition significantly attenuated seizure-like behaviors (e.g., reduced small burst movements) at seizure-inducing concentrations of PTZ. This attenuation was concentration dependent, since higher concentrations of the mTor inhibitor, torin1, exacerbated response to PTZ, suggesting that excessive manipulation of mTor in the context of *hcfc1a* mutation can lead to intractable seizure phenotypes in the zebrafish. These findings align with an established role of mTor hyperactivation in epilepsy and underscore its partial contribution to seizure severity in *hcfc1a*-related pathology.

### Integration with existing knowledge

The mTOR pathway is a well-documented regulator of synaptic plasticity, neuronal excitability, and epileptogenesis [30,31]. Our results extend this paradigm to *HCFC1*-associated disorders, where mTOR dysregulation may arise indirectly via upstream signaling defects, such as AKT hyperphosphorylation. Prior work linking *HCFC1* mutations to GABAergic deficits in mice [25] and radial glial abnormalities in zebrafish [8,12] suggests a multifaceted mechanism, wherein mTOR may underly neurodevelopmental phenotypes and potentially intractable epilepsy in *cblX* syndrome and other HCFC1-related seizure disorders. We observed that mTOR inhibition partially mitigates seizure severity when provided at the appropriate dose, which parallels studies in tuberous sclerosis complex (TSC), where mTOR-targeted therapies reduce but do not eliminate seizures [13–16]. This partial efficacy highlights the likelihood of parallel pathways contributing to HCFC1-related epileptogenesis. Our results also reveal a dose specific response, which may allude to the refractory nature of seizure phenotypes in *cblX* syndrome and may suggest additional pathways at play in *cblX* syndrome and related disorders. Our results warrant the testing of specific doses of mTOR inhibitors alongside traditional seizure treatments (i.e. valproic acid) in *cblX* and related HCFC1 disorders.

### Mechanistic Implications

The blunted PTZ response in *hcfc1a* mutants, combined with persistent pS6 phosphorylation despite torin1 treatment (250nM), suggests intrinsic mTOR pathway dysregulation. This resilience may stem from compensatory feedback mechanisms or crosstalk with other HCFC1-dependent pathways, such as THAP11/ZNF143-mediated transcription [1,30]. Additionally, the light-specific hypomotility reported in *hcfc1a* mutants [8] implies sensory-context-dependent phenotypes, emphasizing the need to evaluate seizures under diverse stimuli. The partial rescue of seizure severity by torin1 aligns with the role of mTOR in regulating neuronal excitability and glial function, both implicated in PTZ responses [27,28].

### Limitations and Technical Considerations

Unexpectedly, vehicle (DMSO) treatment reduced pS6 levels in wildtype larvae, complicating direct comparisons between untreated and torin1 groups. This underscores the importance of vehicle controls in pharmacological studies. Furthermore, our focus on PTZ, a GABA receptor antagonist, may overlook seizure mechanisms specific to other neurotransmitter systems. Future studies using alternative proconvulsants (e.g., kainic acid) could clarify whether mTOR’s role is stimulus dependent. Additionally, the use of larval zebrafish, while advantageous for high-throughput screening, limits investigation of chronic epilepsy models or cognitive comorbidities.

### Therapeutic Relevance

Our findings support precise dose dependent mTOR inhibition as a potential therapeutic strategy for mitigating seizure severity in *cblX* syndrome. However, the incomplete rescue by torin1 suggests combinatorial therapies targeting both mTOR and ancillary pathways (e.g., GABAergic signaling or epigenetic regulators like ASXL1 [10]) may be necessary. Clinical translation will require careful evaluation of mTOR inhibitors in HCFC1 patient-derived models, given the pathway’s pleiotropic roles in neurodevelopment.

### Future Directions

Building on these findings, several avenues merit exploration to deepen our understanding of HCFC1-related epileptogenesis and refine therapeutic strategies. First, the mechanistic dissection of the interplay between HCFC1, mTOR, and downstream effectors could clarify how transcriptional dysregulation converges on neuronal hyperexcitability. Integrating multi-omics approaches such as single-cell RNA sequencing and phosphoproteomics in *hcfc1a* mutants may identify novel mTOR-independent pathways contributing to seizure phenotypes. For instance, the interaction between HCFC1 with epigenetic regulators like ASXL1 [10] and transcriptional complexes such as THAP11/ZNF143 [30] warrants investigation, as these interactions could modulate synaptic gene networks or glial-neuronal crosstalk. Parallel studies in patient-derived induced pluripotent stem cell (iPSC) models would further bridge zebrafish findings to human pathophysiology.

Second, expanding seizure paradigms beyond PTZ could uncover stimulus-specific mechanisms. Optogenetic induction of seizures or audiogenic stimuli may reveal sensory-modulated epileptogenesis, particularly given the light-dependent hypomotility observed in *hcfc1a* mutants [8]. Additionally, chronic seizure models in adult zebrafish could assess whether mTOR inhibition alters epilepsy progression or comorbid cognitive deficits, which are hallmarks of *cblX* syndrome. Third, longitudinal studies evaluating mTOR inhibitor safety, dosage, and efficacy across developmental stages are critical. While acute dose dependent torin1 treatment reduced seizure severity in larvae, prolonged inhibition during neurodevelopment at larger doses might exacerbate HCFC1-related deficits, such as radial glial proliferation [12]. Dose- and timing-dependent studies could optimize therapeutic windows while minimizing off-target effects.

Finally, combinatorial therapies targeting mTOR and complementary pathways such as GABAergic signaling or ASXL1-mediated AKT activation [10] should be explored. Given the partial rescue by torin1, synergistic drug screens could identify compounds that enhance mTOR inhibition or bypass HCFC1-dependent metabolic defects. Collaborations with clinical researchers to profile mTOR activity in *cblX* patients would validate zebrafish findings and accelerate translational applications. Together, these efforts promise to unravel the complexity of HCFC1-associated epilepsy and advance personalized treatment paradigms for neurodevelopmental disorders linked to transcriptional co-factor dysfunction.

## Conclusions

In summary, our work positions mTOR as one potential modulator of seizure severity in *hcfc1a*-deficient zebrafish, offering a mechanistic bridge between HCFC1 dysfunction and epilepsy. While therapeutic targeting of mTOR holds promise, the partial rescue observed here underscores the complexity of *cblX* pathophysiology and the need for multifaceted treatment strategies.

## Supplementary Materials

Figure S1: pS6 dosage gradient

## Author Contributions

Conceptualization, Claudia Gil, Victoria Castro and Anita Quintana; Data curation, Claudia Gil and Annalise Gonzales; Formal analysis, Claudia Gil, David Paz, Victoria Castro, Sepiso Masenga and Anita Quintana; Investigation, Claudia Gil, Antentor Hinton Jr and Anita Quintana; Methodology, Claudia Gil, David Paz, Briana Pinales, Victoria Castro, Claire Perucho, Annalise Gonzales and Anita Quintana; Resources, Giulio Francia and Anita Quintana; Software, Briana Pinales; Supervision, Giulio Francia and Anita Quintana; Validation, David Paz and Annalise Gonzales; Writing – original draft, Victoria Castro, Giulio Francia, Antentor Hinton Jr and Anita Quintana; Writing – review & editing, David Paz, Victoria Castro, Giulio Francia, Sepiso Masenga, Antentor Hinton Jr and Anita Quintana.

## Institutional Review Board Statement

All procedures are approved by the Institutional Animal Care and Use Committee at the University of Texas at El Paso, protocol number 811869-5 or University of Texas at Arlington protocol numbers 25.004, 25.005, 25.006, and 25.007. Methods for euthanasia and anesthesia are performed according to guidelines from the 2020 American Veterinary Medical Association guidebook.

## Data Availability Statement

All data and reagents are available upon request from the corresponding author.

## Acknowledgments

The authors would like to thank the members of the Quintana, Hinton, and Francia labs, past and present, for discussions, animal care, genotyping, and general laboratory maintenance.

