## Supplemental Figures for "A zebrafish seizure model of *cblX* syndrome reveals a dose-dependent refractory response to mTor inhibition"

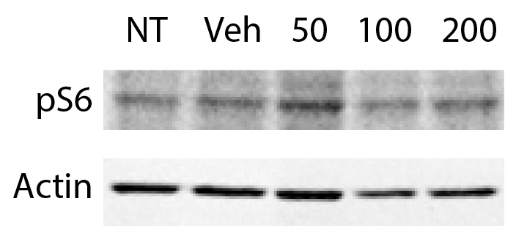

**Figure S1:pS6 dosage gradient.** Western blot was used to determine the optimal concentration of torin1 to reduce mTor activity. Larve were treated as described in the materials and methods and antibodies to detect phosphorylated S6 ribosomal protein and actin were used. NT is non-treated control; veh is vehicle control (DMSO). N=25/group
